# Revisiting rustrela virus – new cases of encephalitis and a solution to the capsid enigma

**DOI:** 10.1101/2021.12.27.474043

**Authors:** Florian Pfaff, Angele Breithaupt, Dennis Rubbenstroth, Sina Nippert, Christina Baumbach, Sascha Gerst, Christoph Langner, Claudia Wylezich, Arnt Ebinger, Dirk Höper, Rainer G. Ulrich, Martin Beer

## Abstract

Rustrela virus (RusV, species *Rubivirus strelense*) is a recently discovered relative of rubella virus (RuV) that has been detected in cases of encephalitis across a wide spectrum of mammals, including placental and marsupial animals. Here we diagnosed two additional cases of fatal RusV-associated meningoencephalitis in a South American coati (*Nasua nasua*) and a Eurasian otter (*Lutra lutra*) that were detected in a zoological garden with history of prior RusV infections. Both animals showed abnormal movement or unusual behaviour and their brains tested positive for RusV using specific RT-qPCR and RNA *in situ* hybridization. As previous sequencing of RusV proved to be very challenging, we employed a sophisticated target-specific capture enrichment with specifically designed RNA baits to generate complete RusV genome sequences from both detected encephalitic animals and apparently healthy wild yellow-necked field mice (*Apodemus flavicollis*). Furthermore, the technique was used to revise three previously published RusV genomes from two encephalitic animals and a wild yellow-necked field mouse. Virus-to-host sequence ratio and thereby sequence coverage improved markedly using the enrichment method as compared to standard procedures. When comparing the newly generated RusV sequences to the previously published RusV genomes, we identified a previously undetected stretch of 309 nucleotides predicted to represent the intergenic region and the sequence encoding the N-terminus of the capsid protein. This indicated that the original RusV sequence was likely incomplete due to misassembly of the genome at a region with an exceptionally high G+C content of >80 mol%, which could not be resolved even by enormous sequencing efforts with standard methods. The updated capsid protein amino acid sequence now resembles those of RuV and ruhugu virus in size and harbours a predicted RNA binding domain that was not encoded in the original RusV genome version. The new sequence data indicate that RusV has the largest overall genome (9,631 nucleotides), intergenic region (290 nucleotides) and capsid protein-encoding sequence (331 codons) within the genus *Rubivirus*.

## Introduction

Rubella virus (RuV; species *Rubivirus rubellae*) was the sole member of the family *Matonaviridae* and the genus *Rubivirus* ^1^, until recently its first relatives rustrela virus (RusV; *Rubivirus strelense*) and ruhugu virus (RuhV; *Rubivirus ruteetense*) were identified ^2^. While RuhV was detected in apparently healthy cyclops leaf-nosed bats (*Hipposideros cyclops*) in Uganda, RusV was associated with cases of fatal neurological disease in placental and marsupial zoo animals in Germany. RusV was initially identified using a metagenomic sequencing workflow from brain tissues of a donkey (*Equus asinus*), a capybara (*Hydrochoeris hydrochaeris*), and a red-necked wallaby (*Macropus rufogriseus)* between July 2018 and October 2019 ^2, 3^. All of these animals were housed in a zoological garden located in northeast Germany, close to the Baltic Sea, and developed acute neurological signs such as ataxia and lethargy, which ultimately resulted in death. RusV was mainly detected in the central nervous system of these animals and only sporadically and in very low concentrations in extraneural organs. RusV-infected wild yellow-necked field mice (*Apodemus flavicollis*) were identified in close proximity to the encephalitic animals’ housings. These rodents were considered as a likely reservoir host, as they carried the virus without obvious encephalitis whereas all tested individuals of other sympatrically occurring rodent species at the same location were RusV-negative ^2^. However, the mode of transmission between potential reservoir and accidental spill-over hosts still remains to be identified ^2^. Currently, no isolates of either RusV or RuhV are available and therefore, most data are limited to *in silico* predictions and analogies with RuV. Furthermore, sequencing of RusV from organ samples proved to be extremely difficult and only three full-length genome sequences and a few partial coding sequences are currently available (**Table 1**).

**Table 1:** Rustrela virus-infected zoo animals, Eurasian otter and yellow-necked field mice from Northern Germany included in this study

| Strain | Organism | Sampling date | Location | Study | Sequence data<sup>#</sup> | Previous accessions | Updated and new full genome accession |
| --- | --- | --- | --- | --- | --- | --- | --- |
| Yellow-necked field mouse/Mu09-1341/2009/Germany | *Apodemus flavicollis* | Jul 2009 | ~2 km distance to zoo | Bennet *et al.* 2020 | T, B | MT274737.1, MT274731.1 | OL960721 |
| Donkey/19_041-1/2019/Germany | *Equus asinus* | Mar 2019 | housed in zoo | Bennet *et al.* 2020 | T, B | MN552442.1 | MN552442.2 |
| Capybara/P19-643/2019/Germany | *Hydrochoerus hydrochaeris* | Oct 2019 | housed in zoo | Bennet *et al.* 2020 | T, B, B+ | MT274724.1 | MT274724.2 |
| Yellow-necked field mouse/KS19-928/2019/Germany | *Apodemus flavicollis* | Sep 2019 | on zoo grounds | Bennet *et al.* 2020 | T, B, B+ | MT274725.1 | MT274725.2 |
| Yellow-necked field mouse/KS20-1296/2020/Germany | *Apodemus flavicollis* | Oct 2020 | ~10 km distance to zoo | Bennet *et al.* 2020 | T, B | MT274732.1, MT274726.1 | OL960722 |
| Yellow-necked field mouse/KS20-1340/2020/Germany | *Apodemus flavicollis* | 2020 | on zoo grounds | Bennet *et al.* 2020 | T, B | MT274733.1, MT274727.1 | OL960726 |
| Yellow-necked field mouse/KS20-1341/2020/Germany | *Apodemus flavicollis* | 2020 | on zoo grounds | Bennet *et al.* 2020 | T, B | MT274734.1, MT274728.1 | OL960725 |
| Yellow-necked field mouse/KS20-1342/2020/Germany | *Apodemus flavicollis* | 2020 | on zoo grounds | Bennet *et al.* 2020 | T, B, B+ | MT274735.1, MT274729.1 | OL960724 |
| Yellow-necked field mouse/KS20-1343/2020/Germany | *Apodemus flavicollis* | 2020 | on zoo grounds | Bennet *et al.* 2020 | T, B, R, P | MT274736.1, MT274730.1 | OL960723 |
| Yellow-necked field mouse/KS20-1512/2020/Germany | *Apodemus flavicollis* | 2020 | on zoo grounds | this study | T, B, R, P | n.a. | OL960720 |
| Yellow-necked field mouse/KS20-1513/2020/Germany | *Apodemus flavicollis* | 2020 | on zoo grounds | this study | T, B | n.a. | OL960719 |
| Yellow-necked field mouse/KS20-1535/2020/Germany | *Apodemus flavicollis* | Jun 2020 | ~10 km distance to zoo | this study | T, B | n.a. | OL960718 |
| South American coati/20_131/2020/Germany | *Nasua nasua* | Aug 2020 | housed in zoo | this study | T, B, B+ | n.a. | OL960717 |
| Eurasian otter/21_002/2020/Germany | *Lutra lutra* | Dec 2020 | ~3 km distance to zoo | this study | T, B, B+ | n.a. | OL960716 |

The genome of rubiviruses consists of single-stranded positive sense (+ss) RNA, that contains two open reading frames (ORF) encoding the non-structural p200 and structural p110 polyproteins, respectively ^4^. Both ORFs are separated by an untranslated intergenic region (IGR). In RuV, co-translational cleavage of the p110 polyprotein results in three structural proteins E1, E2, and the capsid protein ^4^. After cleavage by cellular signal peptidase, the capsid protein remains in the cytoplasm while E1 and E2 enter the secretory pathway using distinct translocation signals ^5^. Based on sequence comparison with RuV, the genomes of RusV and RuhV are likewise predicted to encode the p110 polyprotein and the mature capsid, E1 and E2 proteins ^2^. In RuV, the capsid protein consists of a structurally disordered N-terminal part that contains a RNA-binding domain (RBD) ^6^ and a structurally ordered C-terminal domain (CTD) ^7, 8^ containing the E2 signal sequence ^9^. While the predicted capsid protein sequence and structure of RuhV is analogous to that of RuV, the capsid protein of RusV was considered enigmatic as it appears truncated and lacking e.g. the RBD ^10^.

Here we analysed the RusV genome sequences from two novel cases of RusV encephalitis using a sophisticated target-specific capture enrichment with RNA baits prior to sequencing. This resulted in markedly improved virus-to-background sequence ratios and higher genome coverage particularly in regions of exceptionally high G+C ratios of >80 mol%. The *de novo* assembled sequences suggested a 309 nucleotide (nt) longer RusV genome sequence than initially reported. We also confirmed the sequence extension by reanalysing samples from previously published diseased animals and potential reservoir animals using the same methods, and finally solved the enigma of the unusual RusV capsid protein sequence. So far, the clinical and pathological data are limited to the first description of RusV fatal encephalitis ^2^. We now present further clinical data and an in depth pathological and histopathological evaluation of the two new cases.

## Material and Methods

### Animals and samples included in this study

Brain samples were collected from a South American coati that was housed in a zoological garden in the Northern Germany, a wild Eurasian otter that was found nearby the zoo and three yellow-necked field mice that had been trapped during pest control measures at the zoo (**Table 1**). In addition, samples from previously published animals, including a donkey, a capybara and seven yellow-necked field mice, were re-analysed during this study ^2^.

### Histopathology, immunohistochemistry and RusV RNA *in situ* hybridization

Routine staining, immunohistochemistry (IHC) as well as RNA *in situ* hybridization (RNA ISH) was applied as described earlier with minimal adaptations summarized in Supplemental Table S1 (see also ^2^). Briefly, formalin-fixed, paraffin-embedded (FFPE) brain tissues were processed for haematoxylin and eosin (HE) staining and examination using light microscopy. On consecutive slides, conventional Prussian Blue staining was used to demonstrate haemosiderin, whereas Luxol Fast Blue Cresyl Violet was applied for detection of myelin sheaths and Nissl substance. Immunohistochemistry was performed according to standardized procedures using markers to detect T-cells (CD3), B-cells (CD79a), microglial cells and macrophages (IBA1), astrocytes (glial fibrillary acidic protein, GFAP) and apoptotic cells (active caspase 3). A bright red chromogen labelling was produced with 3-amino-9-ethylcarbazole substrate (AEC, DAKO). Sections were counterstained with Mayer’s haematoxylin. RNA ISH was performed with the RNAScope 2-5 HD Reagent Kit-Red (Advanced Cell Diagnostics, USA) according to the manufacturer’s instructions using a custom-designed probe against the RusV non-structural protein (p200, NSP) ORF, and a negative control probe against the dihydrodipicolinate reductase (*DapB*) gene. Analysis and interpretation were performed by a board-certified pathologist (AB).

### Total RNA extraction for sequencing

Total RNA was extracted from frozen brain tissues as described previously ^11^. Initially, approximately 20 - 30 mg of tissue was snap-frozen in liquid nitrogen and disintegrated using a cryoPREP impactor (Covaris, UK). The pulverized tissue was solubilized in pre-heated lysis buffer AL and RNA was extracted using the RNAdvance Tissue Kit (Beckman Coulter, Germany) in combination with a KingFisher Flex Purification System (Thermo Fisher Scientific, Germany).

### RusV-specific RT-qPCR

RusV-specific RNA was detected by TaqMan RT-qPCR using the AgPath-ID One-Step RT-PCR reagents (Thermo Fisher Scientific, Germany) along with a modified primer/probe set targeting the p200 ORF ^2^. Briefly, 2.5 μl extracted RNA was reverse-transcribed and amplified in a reaction mix of 12.5 μl total volume containing primers RusV_1072_A+ (5’-CGAGCGYGTCTACAAGTTYA-3’; final concentration 0.8 μM) and RusV_1237- (5’-GACCATGATGTTGGCGAGG-3’; 0.8 μM) and probe RusV_1116_A_P (5’-[FAM]CCGAGGARGACGCCCTGTGC[BHQ1]-3’; 0.4 μM). The reaction was performed with the following cycler setup: 45°C for 10 min, 95°C for 10 min, 45 cycles of 95°C for 15 sec, 60°C for 30 sec and 72°C for 30 sec on a Bio-Rad CFX96 qPCR cycler (Bio-Rad, Germany).

### Sequencing of total RNA0

Extracted total RNA was sequenced using a universal metagenomics sequencing workflow ^11, 12^. An amount of 350 ng total RNA per sample was reverse-transcribed into cDNA using the SuperScript IV First-Strand cDNA Synthesis System (Invitrogen, Germany) and the NEBNext Ultra II Non-Directional RNA Second Strand Synthesis Module (New England Biolabs, Germany). Afterwards, cDNA was processed to generate Ion Torrent compatible barcoded sequencing libraries as detailed described ^2, 11^. Libraries were quantified with the QIAseq Library Quant Assay Kit (Qiagen, Germany) and subsequently sequenced on an Ion Torrent S5XL instrument using Ion 530 chips and chemistry for 400 base pair reads (Thermo Fisher Scientific, Germany).

### Sequencing of rRNA-depleted and poly(A)+ enriched RNA

For rRNA depletion we used the NEBNext rRNA Depletion Kit for human, mouse, and rat (New England Biolabs, USA) that specifically depletes cytoplasmic (5S, 5.8S, 18S and 28S rRNA) and mitochondrial ribosomal RNA (12S and 16S rRNA). As the depletion is rRNA sequence-specific, we first confirmed that the human-, mouse- and rat-specific panel would be compatible with samples from yellow-necked field mice by comparing available cytoplasmic and mitochondrial rRNA sequences of all species. Subsequently, 3 μg of the total RNA from two selected yellow-necked field mice were treated with the NEBNext rRNA Depletion Kit for human, mouse, and rat (New England Biolabs), following the manufactures instructions.

Enrichment of poly(A)+ RNA from total RNA was considered appropriate, as the RusV genome, like RuV ^13^, comprises a poly(A) tail at the 3’ terminus. For poly(A)+ enrichment, 3 μg of total RNA from the same yellow-necked field mice were treated with the Dynabeads mRNA DIRECT Micro Purification Kit (Invitrogen, USA) following the manufacturer’s instructions.

Both, rRNA depleted and poly(A)+ enriched RNA, were used for strand-specific library construction with the Collibri Stranded RNA Library Prep Kit (Thermo Fisher, USA). Libraries were quality-checked using a 4150 TapeStation System (Agilent Technologies, USA) with the High Sensitivity D1000 ScreenTape and reagents (Agilent Technologies) and were then quantified using a Qubit Fluorometer (Thermo Fisher) along with the dsDNA HS Assay Kit (Thermo Fisher). Libraries were pooled and sequenced on a NextSeq 500 (Illumina, USA) using a NextSeq 500/550 Mid-output Kit v2.5 with 300 cycles (Illumina).

### Design of custom panRubi bait panels

All available whole-genome sequences of the genus *Rubivirus* were received from NCBI GenBank (86 RuV, one RuhV and three RusV sequences). The genome set was sent to Daicel Arbor Biosciences (Ann Arbor, USA) and a tailored custom myBaits panel for target enrichment via hybridization-based capture was designed. The resulting “panRubi” panel consists of 19,178 RNA oligonucleotide baits with a length of 60 nt arranged every 20 nt along the genomes (designated “panRubi bait set v1”). The set was later supplemented with 22 additional baits covering the newly identified part of the capsid protein-encoding sequence and IGR arranged every 16 nt. This set was mixed with the “panRubi bait set v1” at a ratio of 1:10 to give the “panRubi bait set v2”. All bait sets were checked using Basic Local Alignment Search Tool (BLAST) search against human, mouse, horse, and opossum genomes and no BLAST hit was found.

### Application of RNA baits and sequencing

The custom panRubi bait sets v1/v2 were applied to the sequencing libraries according to the manufacturer’s instructions (myBaits manual v.5.00, Arbor Biosciences, Sep 2020). Hybridization reactions were performed in 1.5 μl safe-lock tubes overlaid with one volume of mineral oil (Carl Roth, Germany), to keep the volume constant during hybridization using a ThermoMixer (Eppendorf, Germany) with 550 rotations per minute. We used the standard protocol (according to ^14^) with a hybridization temperature of 65°C and a hybridization time of about 24 hours. The enriched and purified samples were amplified using the GeneRead DNA Library L amplification Kit (Qiagen, Germany) according to manufacturer’s instructions with 14 cycles and amplicons were purified using solid-phase paramagnetic bead technology. Treated libraries were sequenced after quality check using a Bioanalyzer 2100 (Agilent Technologies) and quantification as described above.

### Read processing and *de novo* assembly

Ion Torrent-derived reads from the myBaits capture enrichment approach were initially quality-trimmed and specific adapters were removed using the 454 Sequencing Systems Software (version 3.0). Instead of host/background removal using specific reference sequences, a G+C content filter was applied to the trimmed reads, as the RusV genome has a particularly high average G+C content of 70.6 mol% ^2^. In detail, only reads with an average G+C content of ≥60 mol% were filtered using PRINSEQ-lite (version 0.20.4; ^15^) and subsequently used for *de novo* assembly with SPAdes genome assembler (version 3.15.2; ^16^) running in single cell mode (--sc) for Ion Torrent data (--iontorrent). The resulting contigs were mapped to the RusV reference sequence MN552442 using Geneious generic mapper (Geneious Prime 2021.0.1) with medium sensitivity allowing discovery of structural variants, and short insertions/deletions (indels) of any size. A consensus sequence was generated and reads were finally mapped back to the consensus sequence using Geneious generic mapper in order to manually inspect genomic termini and possible frameshifts caused by homopolymers.

Illumina-derived reads from rRNA-depleted and poly(A)+-enriched RNA were initially trimmed using Trim Galore (version 0.6.6; ^17^) with automated adapter selection and reads containing only poly(A) homopolymers were trimmed using BBMap/BBDuk (version 38.18, ^18^). For coverage analysis, the trimmed reads of each sample were mapped to the respective assembled genome using Geneious Prime generic mapper in “Low Sensitivity / Fastest” mode. The indexed BAM files were then processed with SAMtools depth (version 1.11; ^19^).

### Phylogenetic analysis

Complete RusV genome sequences were aligned using MAFFT (version 7.450; ^20^) and then used as input for approximately-maximum-likelihood reconstruction with Fast Tree (version 2.1.11; ^21^) using the generalized time-reversible (GTR) model with 5 rate categories and optimized Gamma20 likelihood. The resulting tree was inspected using Geneious Prime (version 2021.0.1).

## Results

### Two carnivoran mammals with neurological disorder

In August 2020, a South American coati (*Nasua nasua*) kept in the zoo showed lethargy, hind limb weakness, convulsion and tremor. Two days later and finally unmoving, the animal was euthanized. Gross pathology revealed swelling of the liver and hyperkeratosis of the footpads. Initial histopathology identified a non-suppurative meningoencephalitis. Findings in the liver included scattered single cell necrosis of hepatocytes and minimal microvesicular fatty change interpreted to be clinically irrelevant, while the footpad hyperkeratosis was interpreted to be age-related. Standard diagnostic tests were negative for mammalian bornaviruses, canine distemper virus and *Salmonella* spp.

In December 2020, a wild Eurasian otter (*Lutra lutra*) was found in the vicinity of the very same zoological garden, without any reported link to the zoo areal, showing abnormal movement. Prior to capturing, the animal was observed in the open waters of the nearby Baltic Sea coast and then later found on the premises of a local school. The animal was sent for clinical observation to the zoo, presenting in a state of malnourishment but with increased food and water uptake, loss of natural shyness and an abrasion at the head indicating blunt trauma. Abnormal movements were still present until the animal was found dead three days later. Pathological examination confirmed hairless spots at the head with a focal perforation of the skin but otherwise non-specific alterations interpreted to be associated with acute, agonal cardiovascular failure. Initial routine histology identified a non-suppurative meningoencephalitis but no further lesions in other organs. Standard diagnostic tests were negative for mammalian bornaviruses, influenza A virus, canine distemper virus, rabies virus, *Salmonella* spp. and *Toxoplasma gondii*.

### Histopathology confirms RusV-associated encephalitis

In general, follow-up histopathology of the RusV-infected South American coati and Eurasian otter confirmed our results previously reported for the RusV-infected donkey, capybara and wallaby from the same zoo. Associated with a non-suppurative meningoencephalitis (**Fig. 1A**), RNA ISH confirmed the presence of RusV-specific RNA within neuronal cell bodies and their processes in both animals (**Fig. 1B** and **C**). Routine HE staining (**Fig. 1A**) as well as Luxol fast blue Cresyl violet staining (Supplemental Fig. S1A) identified neuronal degeneration in the brain of the Eurasian otter but not the South American coati. Scattered cells, in particular perivascularly, were active caspase 3-labelled, indicating subtle apoptosis induction (Supplemental Fig. S1B). Multifocal perivascular cells in brain samples from the otter were positive for iron in the Prussian Blue reaction, confirming intravital hemorrhages (Supplemental Fig. S1C), potentially associated with a suspected history of a blunt trauma. The non-suppurative meningoencephalitis was characterized by perivascular and disseminated infiltrates and few microglial nodules. Immunohistochemistry identified mainly infiltrating CD3-positive T-cells (Supplemental Fig. S1D) but only single CD79-labelled B-cells (Supplemental Fig. S1D inlay). Numerous IBA1-positive microglial cells and infiltrating macrophages were detected intralesionally (Supplemental Fig. S1E). In addition, GFAP immunohistochemistry indicated activation of astrocytes, exhibiting a plump cell shape (Supplemental Fig. S1F).

**Fig. 1:**
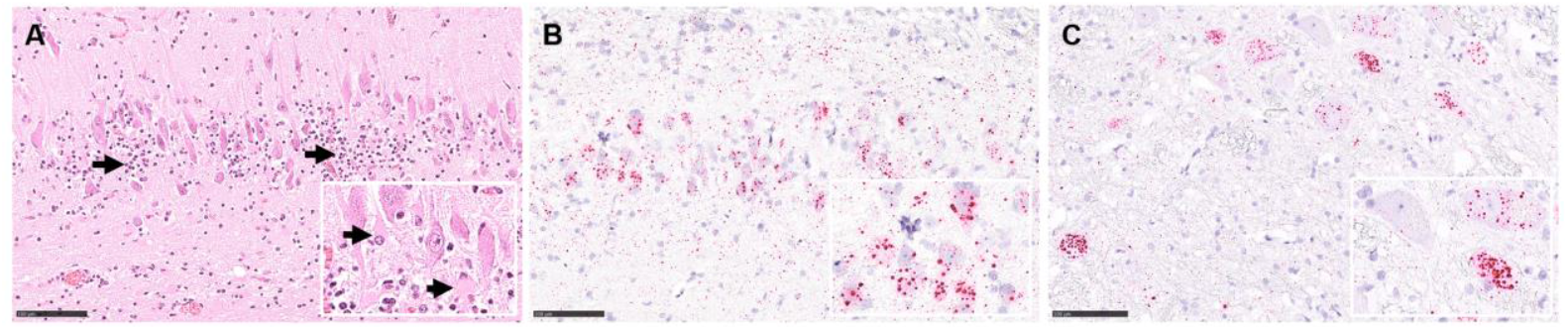
Histopathology from cases of rustrela virus (RusV)-associated meningoencephalitis in a Eurasian otter (*Lutra lutra*) and a South American coati (*Nasua nasua*). (**A**) Non-suppurative meningoencephalitis in the hippocampus region of the otter, with mononuclear infiltrates (arrows) and loss of Nissl substance indicating neuronal degeneration (inlay with arrows), HE stain. Detection of RusV RNA in neurons of the hippocampus region of the otter (**B** and inlay) and brain stem of the coati (**C** and inlay). RNA ISH, chromogenic labelling (fast red) with probes to RusV non-structural polyprotein encoding region, Mayer’s hematoxylin counter stain. Scale bar 100 μm.

### RT-qPCR confirms presence of RusV in encephalitic animals and reservoir hosts

A RusV-specific RT-qPCR confirmed the presence of viral RNA in the brain of the South American coati (Cq 18.9) and the Eurasian otter (Cq 22.5). Furthermore, using the same RT-qPCR setup, we also reanalysed samples from two previously investigated zoo animals and from ten previously published or recently collected RusV-infected wild yellow-necked field mice from within and around the zoo (**Table 1**). Cq values of frozen brain samples ranged from 15.1 to 25.8, with a median of 18.4 (**Fig. 2A**), whereas FFPE brain tissue from a capybara with encephalitis revealed the highest Cq of 27.6 corresponding to the lowest amount of detectable RNA.

**Fig. 2:**
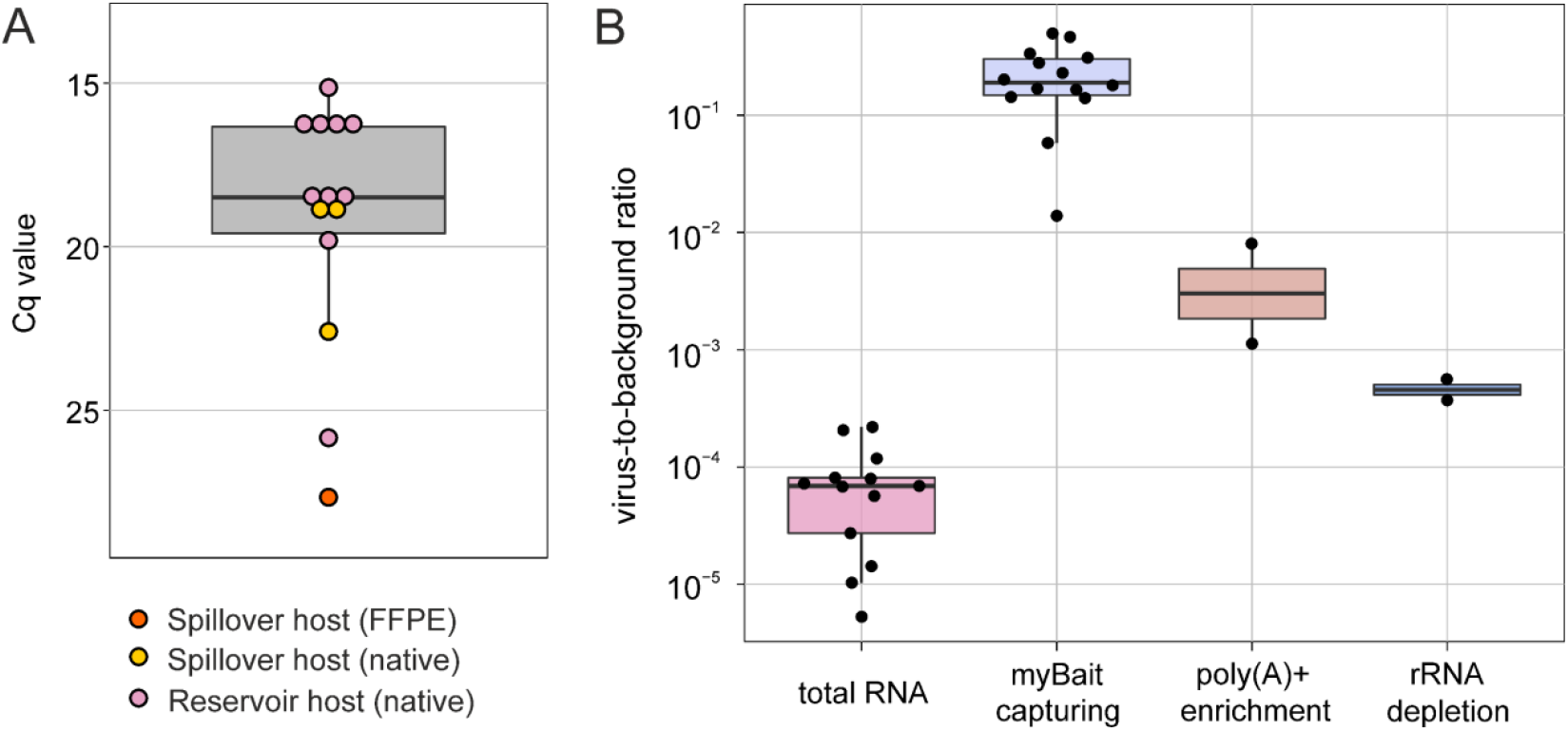
**(A)** Comparison of RusV-specific RT-qPCR Cq values for frozen or FFPE brain samples from potential reservoir (yellow-necked field mouse) and spill-over hosts (donkey, capybara, South American coati, European otter). **(B)** Comparison of virus-to-background sequence ratio observed in sequencing data using different RNA preparations and post library capturing methods.

### Increasing RusV sequencing efficiency

Initially, we used total RNA from brain samples of the Eurasian otter and South American coati for sequencing and *de novo* assembly. However, this resulted in incomplete and highly fragmented genome sequences due to very low virus-to-background sequence ratios of 0.021% and 0.001% for the South American coati and Eurasian otter, respectively. The virus-to-background ratios observed during sequencing of total RNA from all samples included in this study (**Table 1**) ranged from 0.00053% to 0.022%, with a median of 0.0069% (**Fig. 2B**). To increase the efficiency of RusV sequencing, we compared poly(A)+ enrichment, rRNA depletion as well as post library hybridization-based capturing (bait capturing) for selected samples.

By reducing host-derived RNA, poly(A)+ enrichment and rRNA depletion increased virus-to-background sequence ratios by factors of 67 and 6.8, respectively, as compared to total RNA, resulting in median virus-to-background sequence ratios of 0.46% and 0.047%, respectively (**Fig. 2B**). The application of bait capturing to libraries prepared from total RNA achieved virus-to-background sequence ratios of 1.4% to 49.9% with a median of 19.1% (**Fig. 2B**), corresponding to a median 2,772-fold increase.

The characteristic sequence coverage pattern observed for bait-captured libraries closely resembled that of total RNA sequencing (Supplemental Fig. S2). In contrast, libraries from poly(A)+ enriched RNA had a strong bias in coverage towards the 3’ end of the RusV genome. Depletion of rRNA resulted in a relatively uniform coverage across the genome with a bias towards the 5’ end of the genome (Supplemental Fig. S2). No coverage dropout was noted for any of the applied methods.

### Generation and comparison of full length RusV genomes

As bait capturing proved to be very efficient, we applied the technique to all 14 available brain samples (**Table 1**), including South American coati and Eurasian otter, and used the sequencing data for *de novo* assembly. The assembly of each library resulted in contigs that were matched to the RusV genome MN552442. For all samples, a full-length RusV genome without any gaps could be derived from the matched *de novo* assembled contigs.

An alignment of all 14 RusV genome sequences showed only minor variation with an overall pairwise nt identity of 98.8%. Phylogeny based on the aligned whole genome sequences confirmed the high genetic identity of the RusV genomes originating from within or in close proximity of the zoo (Supplemental Fig. S3). RusV sequences from apparently healthy yellow-necked field mice and encephalitic mammals, including the South American coati and Eurasian otter, clustered closely together. Notably, two RusV sequences from yellow-necked field mice collected in a distance of about 10 km from the zoo grouped in a separate genetic branch (Supplemental Fig. S3).

### Revised RusV genome and implications for the IGR and capsid protein-coding sequence

All 14 RusV genomes assembled from bait-captured libraries showed a 309 nt stretch ranging from pos. 6,062 to pos. 6,370 and covering part of the IGR and the N-terminal part of the capsid protein-coding sequence. This stretch had not been present in the three initially released RusV genomes generated by total RNA sequencing ^2^ but was now identified when re-sequencing the very same sample materials using bait capturing (**Fig. 3A**). As the panRubi v1 bait set did not comprise the extra 309 nt-long region, the bait set was complemented with probes specifically targeting this region, leading to a further improved coverage within the respective region (Supplemental Fig. S4).

**Fig. 3:**
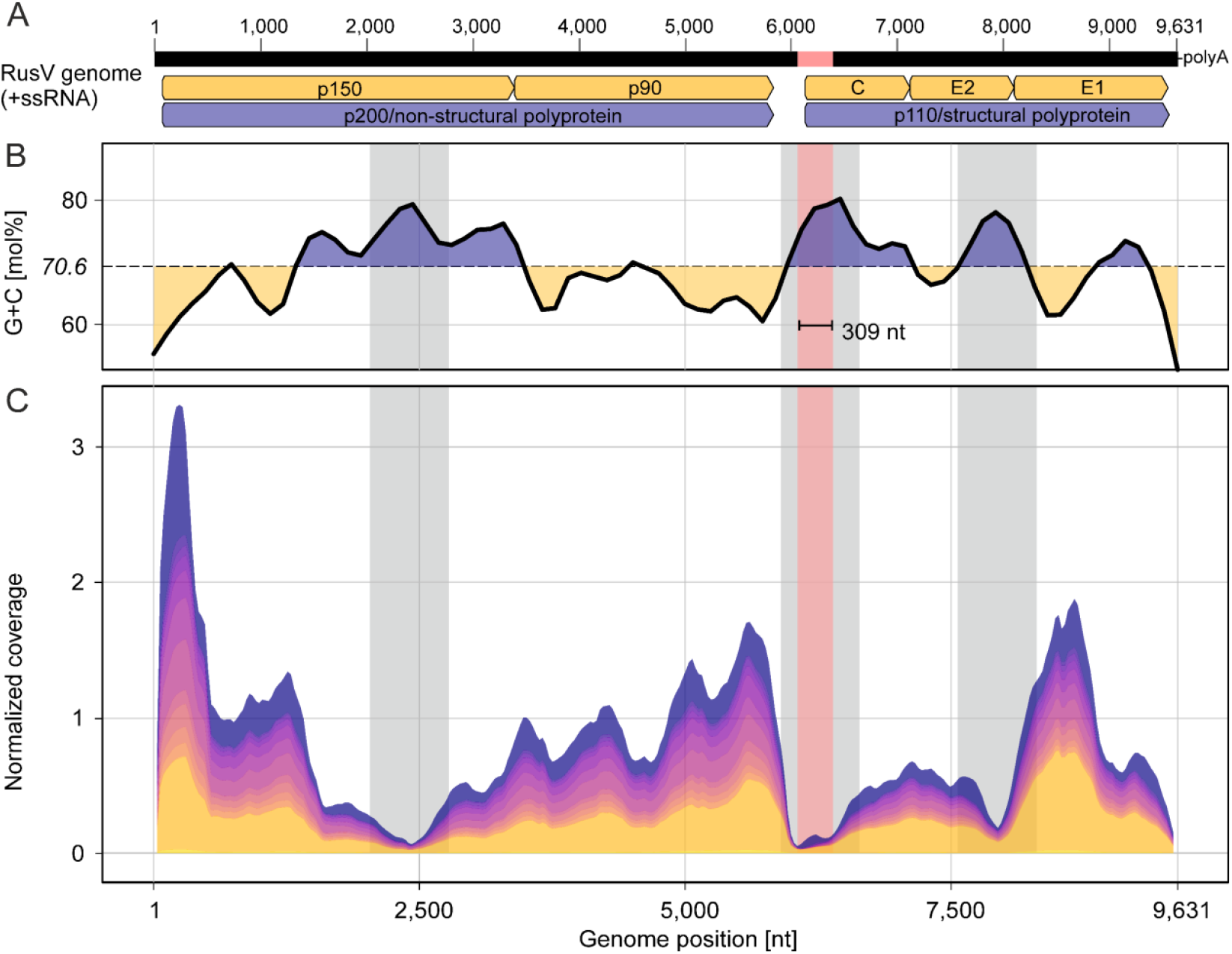
Schematic rustrela virus (RusV) genome sequence **(A)** showing averaged G+C content **(B)** and cumulated RusV sequence coverage of all 14 animals included in this study **(C)**. The newly identified 309 nt sequence stretch partly covering the intergenic region and p110 ORF is highlighted in red. Note that the start of the p110 coding ORF is located within the newly identified sequence stretch, leading to a longer capsid protein sequence as compared to the previously published RusV genomes. Grey labelled areas in **B** and **C** indicate areas of particularly high G+C content.

In general, the observed sequencing coverage varied considerably across the genome, showing pronounced maxima and minima in all samples (**Fig. 3C**). The three genomic regions with the most prominent reduction in sequence coverage correlated with the highest G+C content (**Fig. 3B** and **C**), while genome regions with very high coverage correlated with lower G+C content. The newly identified 309 nt region correlated with a G+C peak and possessed a particularly low sequence coverage (**Fig. 3A-C**).

As a consequence, the IGR and the predicted p110 ORF are longer than initially reported (**Fig. 4A** and **B**). The IGR of all 14 full-length RusV sequences is spanning 290 nt between the stop codon of the predicted p200 ORF and start codon of the p110 ORF. In comparison, the IGRs of RuhV and RuV were reported to be 75 nt and 120 nt in length, respectively (**Fig. 4A**). Based on an ATG start codon in the newly identified 309 nt region and a prediction of the signal peptidase cleavage site (Supplemental Fig. S5), the predicted capsid protein-encoding sequence of RusV is 996 nt (332 amino acids, aa) in length. In comparison, the length of the capsid protein-encoding regions of RuhV and RuV was predicted to be 951 nt (317 aa) and 900 nt (300 aa), respectively (**Fig. 4B**).

**Fig. 4:**
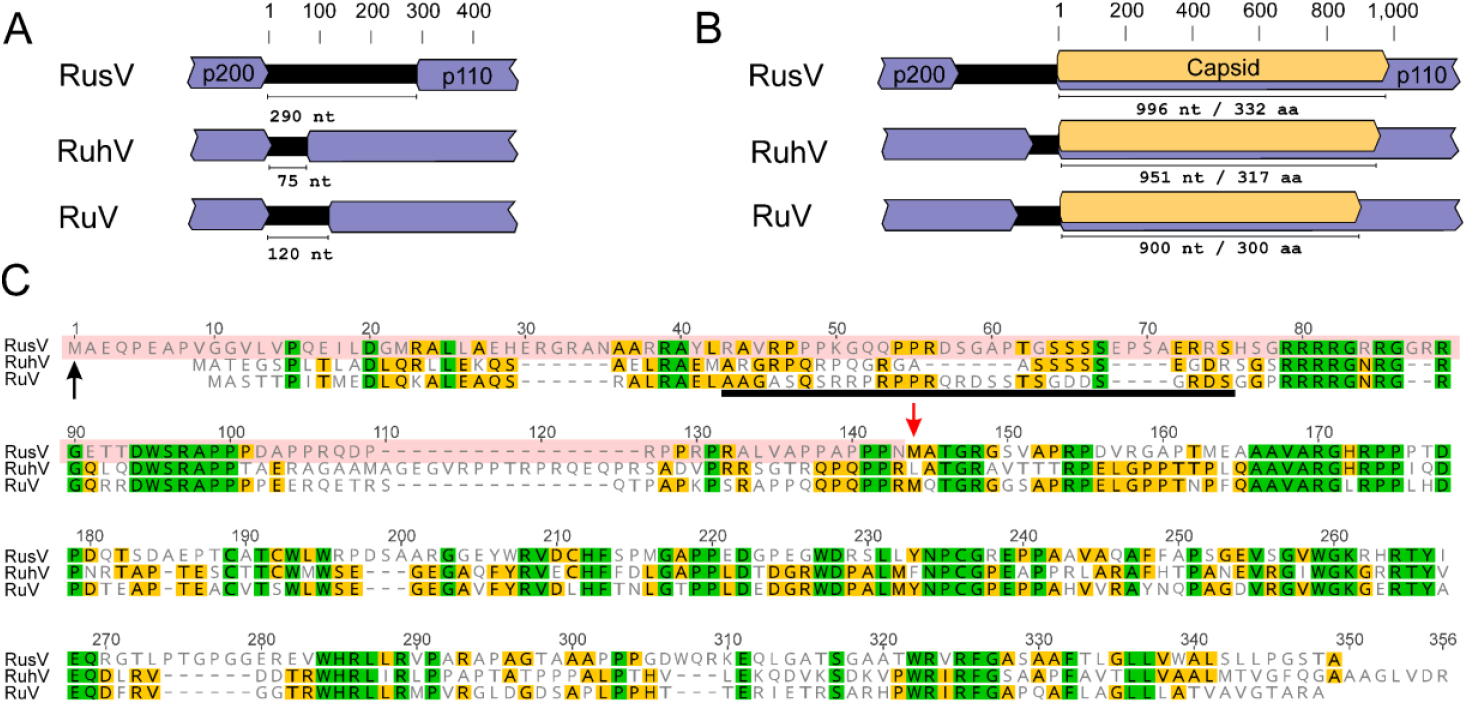
Comparision of the rubivirus intergenic region and capsid protein-encoding sequences. (**A**) The size of the intergenic region between the non-structural (p200) and structural polyprotein (p110) ORFs of rustrela virus (RusV; MN552442.2), ruhugu virus (RuhV; MN547623) and rubella virus (RuV; NC_001545) is shown. (**B**) The predicted length of the capsid protein-coding sequence (highlighted in orange) is shown for RusV, RuhV and RuV. (**C**) The sequences of the capsid protein from RusV, RuhV and RuV are compared using an amino acid sequence alignment. Amino acid residues highlighted in green or yellow are conserved in all three or at least in two of the viruses, respectivly. The N-terminal part of the RusV mature capsid protein (highlighted in red; start marked by black arrow) has been determined in this study. The red arrow indicates the predicted start of the capsid protein in the previously published RusV sequence. The RNA binding site of the RuV capsid protein is indicated by the black bar.

Comparison of the revised capsid protein sequence of RusV to both RuhV and RuV revealed highly conserved stretches (**Fig. 4C**). The revised RusV capsid protein sequence comprised a part that has been predicted to be the RBD in RuV. This part had been absent in the initially published RusV genome. Directly downstream of the predicted RuV RBD region, a polybasic motif (RRRRG R/N RG) can be found that is highly conserved between RusV, RuhV and RuV. This polybasic motif is followed by a likewise highly conserved hydrophobic motif (DWSRAPP).

## Discussion

We recently discovered RusV in the central nervous system of encephalitic zoo animals and wild yellow-necked field mice on the basis of metagenomic sequencing of total RNA, RT-qPCR and RNA ISH ^2^. More than 300 million reads from different sequencing platforms and numerous samples and subsamples were used in order to generate the first RusV genomes originating from three individuals (MN552442.1, MT274724.1 and MT274725.1). A combination of *de novo* assembly, mapping, BLASTx, and manual inspection was used to generate these RusV genomes ^2^. The assembly was exceptionally difficult, as most parts of the RusV genome have a G+C content of >70 mol% with low complexity G+C stretches and coverage dropping drastically at several positions (**Fig. 3B** and **C**). In the IGR, the G+C content even exceeds 85 mol%. Nevertheless, despite very large efforts in sequence determination and characterisation of the RusV genome, questions remained regarding its unusual IGR and capsid protein, which appeared to be rather short in comparison to RuV and RusV and lacked a potential RBD ^10^.

We now investigated two new cases of RusV-associated fatal meningoencephalitis in a European otter and a South American coati that clinically and histologically closely resembled previous RusV-associated cases and that further broadened the spectrum of infected mammals, which now includes placental mammals of the orders Rodentia (families Caviidae and Muridae), Carnivora (families Procynoidae and Mustelidae), and Perissodactyla (family Equidae) as well as marsupials of the order Diprotodontia (family Macropodidae). This broad host spectrum is in clear contrast to rubella virus, for which humans are the only host ^8^. So far, we assume that the wild yellow-necked field mouse may act as reservoir host. However, the transmission route to the other hosts remains unclear.

Despite the relatively low RusV-specific RT-qPCR Cq values in native organ samples, detected using a RusV-specific RT-qPCR (**Fig. 2A**), the virus-to-host sequence ratio observed when sequencing total RNA was unsatisfying. It has been shown, that genome length, virus species, virus- and host-derived RNA concentrations as well as the overall composition of the sample matrix impact the virus-background ratio ^22^. RT-qPCR results often do not reflect this complex interplay and may lead to false expectations for sequencing.

As sequencing of RusV genomes from total RNA proved to be very difficult, we attempted to increase the sequencing efficiency by poly(A)+ enrichment, rRNA depletion and post-library bait capturing. Poly(A)+ enrichment was more efficient than rRNA depletion, resulting in higher virus-to-background ratios (**Fig. 2B**). This observation is in in accordance with other studies, in which rRNA-depleted RNA preparations contained many host-derived small or long non-coding RNAs that are absent in poly(A)+ enriched RNA preparations ^23^. However, poly(A)+ selection introduced a 3’ sequence coverage bias resulting in poor 5’ coverage. This bias has been reported previously for poly(A)+ enrichment methods and is most likely caused by partially degraded transcripts particularly in samples with highly degraded RNA ^24^. RNA quality and integrity plays a major role in sequencing experiments and is directly connected to sampling conditions, transportation and storage ^25–27^. While we used qualified and robust methods for RNA extraction and preservation ^11^, the RNA preparations used for comparison of the different methods originated from brain tissues of two wild-trapped rodents that were sampled under suboptimal conditions.

Hybridization-based capturing has been shown to markedly increase efficiency of RNA virus sequencing previously ^14, 28–30^ and was also found to be most efficient in this study, increasing the median virus-to-background sequence ratio 2,772-fold. Using this technique, we sequenced or re-sequenced 14 full-length RusV genomes from cases of encephalitis and from wild yellow-necked field mice. Using the hybridization-based bait capturing method, the overall sequence coverage and especially the coverage in challenging regions was markedly improved. However, we found a correlation between sharp drops in sequencing coverage within regions of very high G+C content exceeding ∼75 mol%. This may indicate a technical limit of the used sequencing platforms and has been described for different technologies ^31–34^. It has also been suggested, that extreme G+C contents may negatively affect *de novo* assemblies ^35^.

Within a region of high G+C content, spanning IGR and the 5’-end of the capsid protein-encoding sequence, we now found a notable sequence difference, namely a previously unidentified stretch of 309 nt, in comparison to the initially reported RusV genome. Thereby, the predicted capsid protein of RusV is longer than described earlier and now includes the typical rubivirus capsid protein features, such as the RBD that might be crucial for virion formation ^6, 7^. The RBD was unexpectedly missing in the initially annotated RusV capsid protein as pointed out recently in detail by Das and Kilian ^10^. An alignment of the capsid proteins of RusV, RuhV and RuV showed highly conserved motifs that were initially absent in the predicted RusV protein. However, the region identified as RBD in RuV ^6^ appears to be only poorly conserved on aa sequence level in the RuhV or RusV when compared in an alignment. Whether conserved motifs are involved in RNA binding or other structural features remains unclear, as no structural model is currently available for the N-terminal part of the RuV capsid protein ^7^.

The revised version of the RusV genome reveals RusV to have the longest capsid protein-encoding sequence and IGR among all three currently known matonaviruses (**Fig. 2A** and **B**). It has been shown for RuV, that the p110 polyprotein is translated from a subgenomic RNA by using a separate promoter within the IGR ^36, 37^. However, based on the coverage along the genome, we could not find any indication for the presence of subgenomic RNA in the analysed samples. This may indicate that either RusV does not translate the p110 polyprotein from a subgenomic RNA or the RusV replication cycle includes stages without presence of subgenomic RNA. Future studies should address these open questions.

In conclusion, we were able to markedly increase RusV sequencing efficiency leading to an improved genome coverage by employing a bait capturing-based enrichment strategy. Overall, 14 high-quality whole-genomes from RusV-related encephalitis cases and reservoir hosts could be generated applying this strategy. By *de novo* assembly, we identified an extra 309 nt sequence spanning the partial RusV IGR and 5’-end of the capsid protein-encoding region. The RusV example impressively demonstrates the difficulties in correctly determining sequences with an extreme G+C content, but also suggests possible solutions that are now available, such as targeted enrichment via RNA baits. The updated RusV sequence now allows further studies about the function of conserved regions of RusV, but also about viral replication using reverse genetics.

## Acknowledgement

We thank Daicel Arbor Biosciences for providing additional baits that quickly allowed complementing the panRubi panel. We thank Patrick Zitzow, Jenny Lorke, Kathrin Steffen, Doreen Schulz, Silvia Schuparis and Gabriele Czerwinski for excellent technical assistance.

## Data availability

Revised versions of previously published RusV genome sequences are available under DDBJ/ENA/GenBank accession numbers MN552442.2, MT274724.2, and MT274725.2. Novel RusV genomes from this study are available under DDBJ/ENA/GenBank accession numbers: OL960716 - OL960726

## Ethics statement

This study involved no animal experiments. All animal materials were from routine diagnostics or pest rodent control measures.

## Author contributions

**Conceptualization:** DR, MB, RGU Data Curation: FP

**Formal analysis:** FP, AB

**Investigation:** FP, AB, DR, SN, CB, SG, CL, CW, AE, DH

**Supervision:** DR, MB, RGU Visualization: FP, AB

**Writing - Original Draft:** FP, AB, DR

**Writing - Review & Editing:** FP, AB, DR, SN, CB, SG, CL, CW, AE, DH, RGU, MB

## References

1. Rubing Chen, Suchetana Mukhopadhyay, Andres Merits, Bethany Bolling, Farooq Nasar, Lark L. Coffey, Ann Powers, Scott C. Weaver, Donald Smith, Peter Simmonds and Stuart Siddell (2018) Create a new family Matonaviridae to include the genus Rubivirus, removed from the family Togaviridae. https://talk.ictvonline.org/ictv/proposals/2018.013S.A.v3.Matonaviridae.zip. Accessed 28 Jul 2021

2. Bennett AJ, Paskey AC, Ebinger A et al. (2020) Relatives of rubella virus in diverse mammals. Nature 586:424–428. https://doi.org/10.1038/s41586-020-2812-9

3. Bennett AJ, Paskey AC, Ebinger A et al. (2020) Author Correction: Relatives of rubella virus in diverse mammals. Nature 588:E2. https://doi.org/10.1038/s41586-020-2897-1

4. Oker-Blom C (1984) The gene order for rubella virus structural proteins is NH2-C-E2-E1-COOH. J Virol 51:354–358

5. Oker-Blom C, Jarvis DL, Summers MD (1990) Translocation and cleavage of rubella virus envelope glycoproteins: identification and role of the E2 signal sequence. J Gen Virol 71 (Pt 12):3047–3053. https://doi.org/10.1099/0022-1317-71-12-3047

6. Liu Z, Yang D, Qiu Z et al. (1996) Identification of domains in rubella virus genomic RNA and capsid protein necessary for specific interaction. J Virol 70:2184–2190

7. Mangala Prasad V, Willows SD, Fokine A et al. (2013) Rubella virus capsid protein structure and its role in virus assembly and infection. Proc Natl Acad Sci U S A 110:20105–20110. https://doi.org/10.1073/pnas.1316681110

8. Das PK, Kielian M (2021) Molecular and Structural Insights into the Life Cycle of Rubella Virus. J Virol. https://doi.org/10.1128/JVI.02349-20

9. Suomalainen M, Garoff H, Baron MD (1990) The E2 signal sequence of rubella virus remains part of the capsid protein and confers membrane association in vitro. J Virol 64:5500–5509. https://doi.org/10.1128/JVI.64.11.5500-5509.1990

10. Das PK, Kielian M (2021) The Enigmatic Capsid Protein of an Encephalitic Rubivirus. J Virol. https://doi.org/10.1128/JVI.02294-20

11. Wylezich C, Papa A, Beer M et al. (2018) A Versatile Sample Processing Workflow for Metagenomic Pathogen Detection. Sci Rep 8. https://doi.org/10.1038/s41598-018-31496-1

12. Forth LF, Höper D (2019) Highly efficient library preparation for Ion Torrent sequencing using Y-adapters. Biotechniques 67:229–237. https://doi.org/10.2144/btn-2019-0035

13. Dominguez G, Wang C-Y, Frey TK (2004) Sequence of the genome RNA of rubella virus: Evidence for genetic rearrangement during togavirus evolution. Virology 177:225–238. https://doi.org/10.1016/0042-6822(90)90476-8

14. Wylezich C, Calvelage S, Schlottau K et al. (2021) Next-generation diagnostics: virus capture facilitates a sensitive viral diagnosis for epizootic and zoonotic pathogens including SARS-CoV-2. Microbiome 9:51. https://doi.org/10.1186/s40168-020-00973-z

15. Schmieder R, Edwards R (2011) Quality control and preprocessing of metagenomic datasets. Bioinformatics 27:863–864. https://doi.org/10.1093/bioinformatics/btr026

16. Bankevich A, Nurk S, Antipov D et al. (2012) SPAdes: a new genome assembly algorithm and its applications to single-cell sequencing. J Comput Biol 19:455–477. https://doi.org/10.1089/cmb.2012.0021

17. Felix Krueger, Frankie James, Phil Ewels et al. (2021) TrimGalore: v0.6.7. https://github.com/FelixKrueger/TrimGalore. Accessed 28 Jul 2021

18. Brian Bushnell BBTools suite. https://jgi.doe.gov/data-and-tools/bbtools/. Accessed 23 Jul 2021

19. Li H, Handsaker B, Wysoker A et al. (2009) The Sequence Alignment/Map format and SAMtools. Bioinformatics 25:2078–2079. https://doi.org/10.1093/bioinformatics/btp352

20. Katoh K, Standley DM (2013) MAFFT Multiple Sequence Alignment Software Version : Improvements in Performance and Usability. Mol Biol Evol 30:772–780. https://doi.org/10.1093/molbev/mst010

21. Price MN, Dehal PS, Arkin AP (2010) FastTree 2 – approximately maximum-likelihood trees for large alignments. PLoS One 5:e9490. https://doi.org/10.1371/journal.pone.0009490

22. Ebinger A, Fischer S, Höper D (2021) A theoretical and generalized approach for the assessment of the sample-specific limit of detection for clinical metagenomics. Comput Struct Biotechnol J 19:732–742. https://doi.org/10.1016/j.csbj.2020.12.040

23. Zhao S, Zhang Y, Gamini R et al. (2018) Evaluation of two main RNA-seq approaches for gene quantification in clinical RNA sequencing: polyA+ selection versus rRNA depletion. Sci Rep 8:4781. https://doi.org/10.1038/s41598-018-23226-4

24. Cui P, Lin Q, Ding F et al. (2010) A comparison between ribo-minus RNA-sequencing and polyA-selected RNA-sequencing. Genomics 96:259–265. https://doi.org/10.1016/j.ygeno.2010.07.010

25. Sidova M, Tomankova S, Abaffy P et al. (2015) Effects of post-mortem and physical degradation on RNA integrity and quality. Biomol Detect Quantif 5:3–9. https://doi.org/10.1016/j.bdq.2015.08.002

26. Bauer M, Gramlich I, Polzin S et al. (2003) Quantification of mRNA degradation as possible indicator of postmortem interval – a pilot study. Leg Med (Tokyo) 5:220–227. https://doi.org/10.1016/j.legalmed.2003.08.001

27. Bonadio RS, Nunes LB, Moretti PNS et al. (2021) Insights into how environment shapes post-mortem RNA transcription in mouse brain. Sci Rep 11:13008. https://doi.org/10.1038/s41598-021-92268-y

28. Metsky HC, Matranga CB, Wohl S et al. (2017) Zika virus evolution and spread in the Americas. Nature 546:411–415. https://doi.org/10.1038/nature22402

29. Piantadosi A, Kanjilal S, Ganesh V et al. (2018) Rapid Detection of Powassan Virus in a Patient With Encephalitis by Metagenomic Sequencing. Clin Infect Dis 66:789–792. https://doi.org/10.1093/cid/cix792

30. Matranga CB, Andersen KG, Winnicki S et al. (2014) Enhanced methods for unbiased deep sequencing of Lassa and Ebola RNA viruses from clinical and biological samples. Genome Biol 15:519. https://doi.org/10.1186/PREACCEPT-1698056557139770

31. Browne PD, Nielsen TK, Kot W et al. (2020) GC bias affects genomic and metagenomic reconstructions, underrepresenting GC-poor organisms. Gigascience 9. https://doi.org/10.1093/gigascience/giaa008

32. Dohm JC, Lottaz C, Borodina T et al. (2008) Substantial biases in ultra-short read data sets from high-throughput DNA sequencing. Nucleic Acids Res 36:e105. https://doi.org/10.1093/nar/gkn425

33. Quail MA, Smith M, Coupland P et al. (2012) A tale of three next generation sequencing platforms: comparison of Ion Torrent, Pacific Biosciences and Illumina MiSeq sequencers. BMC Genomics 13:341. https://doi.org/10.1186/1471-2164-13-341

34. Benjamini Y, Speed TP (2012) Summarizing and correcting the GC content bias in high-throughput sequencing. Nucleic Acids Res 40:e72. https://doi.org/10.1093/nar/gks001

35. Chen Y-C, Liu T, Yu C-H et al. (2013) Effects of GC bias in next-generation-sequencing data on de novo genome assembly. PLoS One 8:e62856. https://doi.org/10.1371/journal.pone.0062856

36. Tzeng W-P, Frey TK (2002) Mapping the rubella virus subgenomic promoter. J Virol 76:3189–3201. https://doi.org/10.1128/JVI.76.7.3189-3201.2002

37. Oker-Blom C, Ulmanen I, Kääriäinen L et al. (1984) Rubella virus 40S genome RNA specifies a 24S subgenomic mRNA that codes for a precursor to structural proteins. J Virol 49:403–408

