## Supplemental data for "Revisiting rustrela virus – new cases of encephalitis and a solution to the capsid enigma"

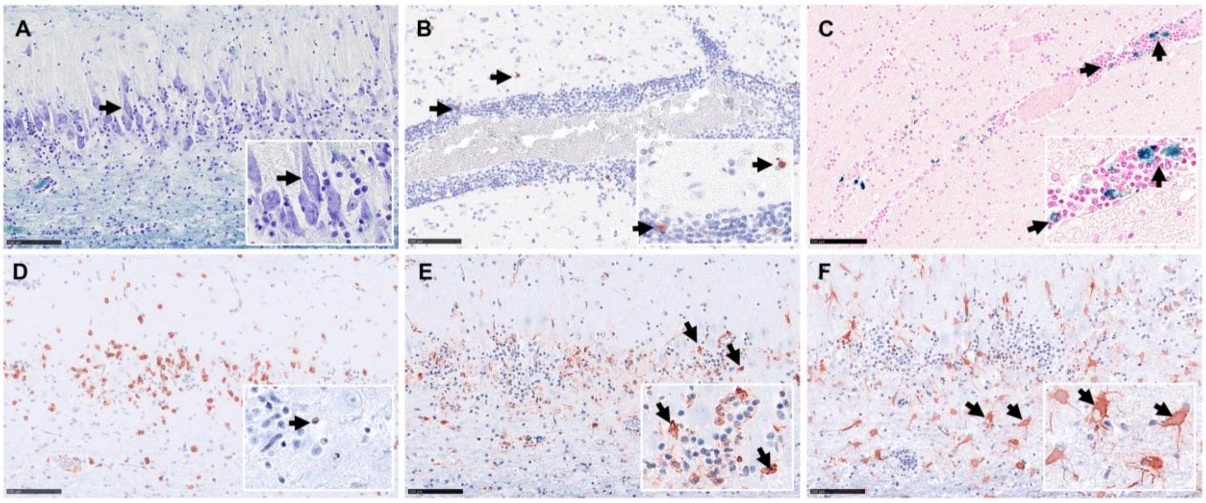


**Fig. S1:** Special stains and immunohistochemistry from cases of rustrela virus (RusV)-induced meningoencephalitis exemplarily shown here for the Eurasian otter in the hippocampus region (**A**, **D-F**) and cortex (**B**, **C**). Loss of Nissl substance with central chromatolysis (arrow) indicating neuronal degeneration, Luxol fast blue cresyl violet stain (**A**, and inlay). Few active caspase 3-immunoreactive (arrows) apoptotic cells (**B**, and inlay). Multifocal perivascular cells, positive for iron in the Prussian Blue reaction (arrows) indicating intravital haemorrhage (**C** and inlay). Numerous infiltrating CD3-labelled T-lymphocytes (**D**) but only single CD79a labelled B-cells in the same region (inlay with arrow). High numbers of plump Iba-1 immunoreactive (arrows) microglial cells and infiltrating macrophages (**E** and inlay). Many GFAP-immunoreactive astrocytes (arrows) with plump cell shape indicating astroglial activation (**F** and inlay). (**B**, **D-F**) immunohistochemistry, AEC chromogen, Mayer's haematoxylin counter stain. All scale bars 100 µm.

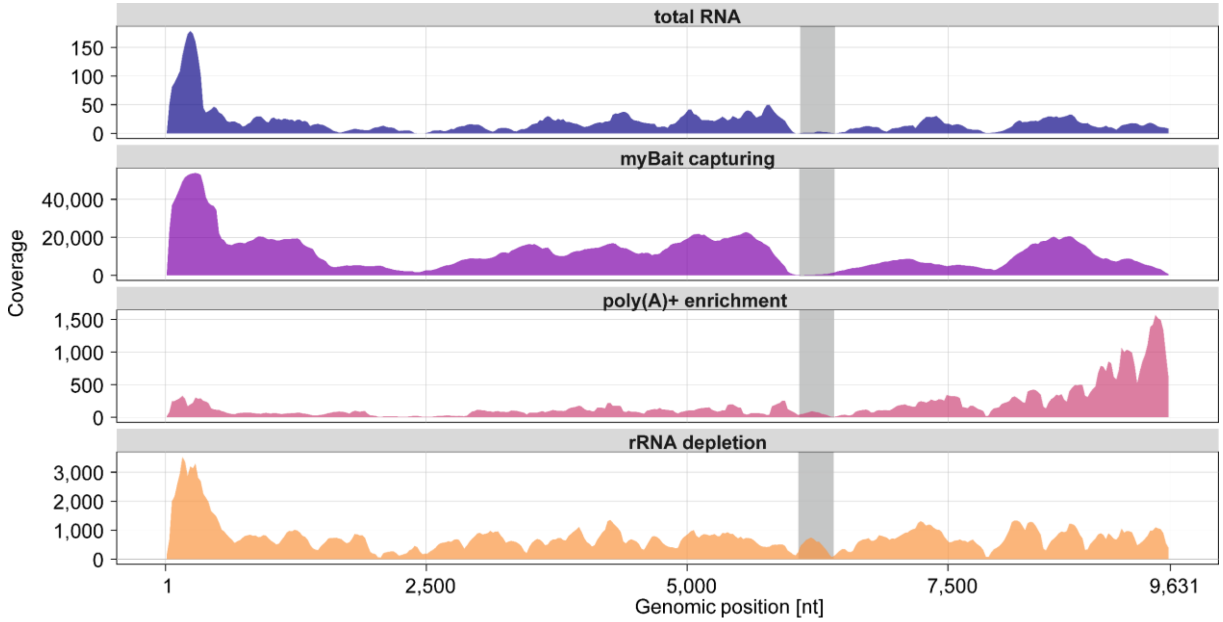


**Fig. S2.** Comparison of sequencing coverage across the rustrela virus (RusV) genome using untreated total RNA and differently enriched or depleted RNA preparations and post library capturing methods. Coverage obtained for the different methods is shown for the sample "Yellow-necked field mouse/KS20-1342/2020/Germany". The newly identified sequence stretch in the intergenic region and capsid protein-encoding sequence is highlighted as grey area.

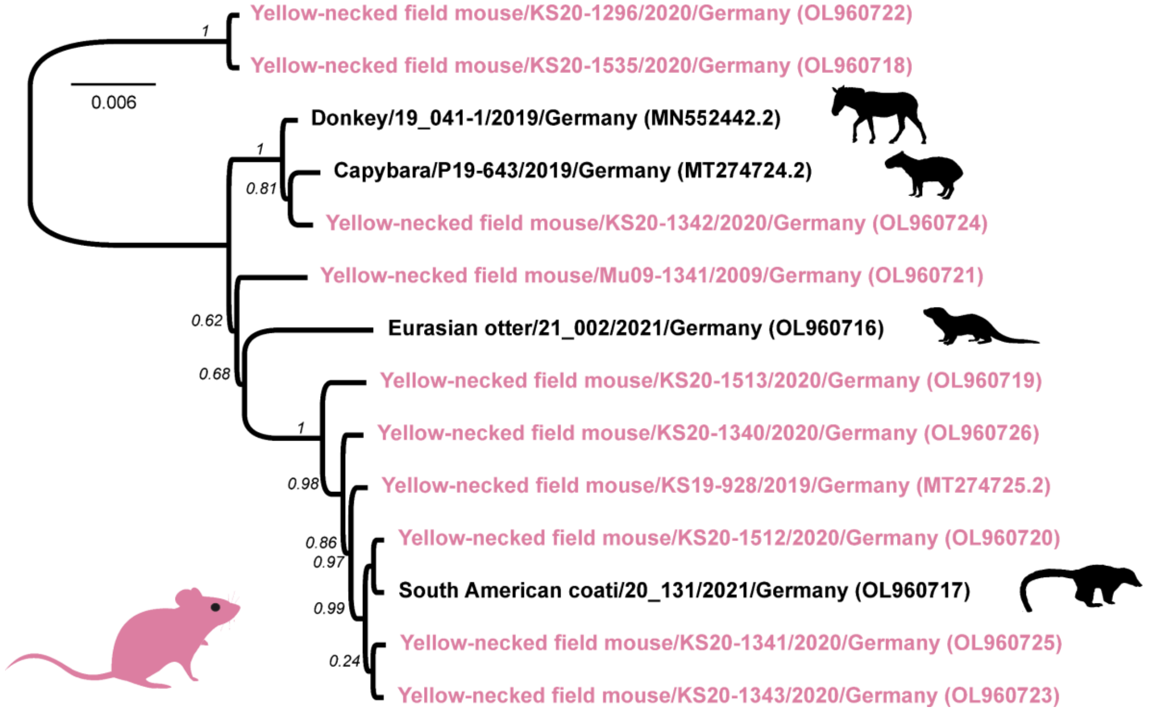


**Fig. S3.** Phylogenetic tree for all available rustrela virus (RusV) full genome sequences. RusV sequences from yellow-necked field mice are highlighted in red while RusV sequences from potential spill-over hosts succumbed to meningoencephalitis are depicted in black. The tree was reconstructed using approximately-maximum-likelihood as implemented in Fast Tree (version 2.1.11; GTR model, 5 rate categories and optimized Gamma20 likelihood). Branch support is indicated as italic numbers.

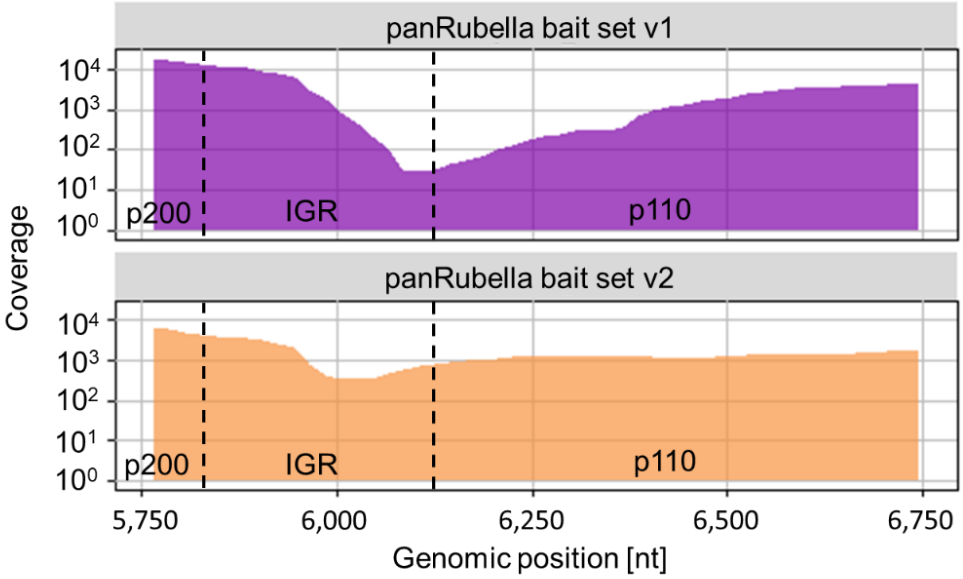


**Fig. S4.** Differences in sequence depth across the intergenic region (IGR) and parts of the non-structural (p200) and structural polyprotein (p110) encoding sequences of rustrela virus (RusV) in high-throughput sequencing using different myBait capturing sets. Coverage is shown for a total RNA library from the sample "Yellow-necked field mouse/KS20-1342/2020/Germany" after treatment with the initial panRubi bait set v1 (violet) or the updated myBait set (panRubi bait set v2; orange) additionally containing baits targeting the newly identified 309 nt stretch partially covering IGR and capsid protein-encoding sequence. The mean coverage is shown using a sliding window (window size 10 nt, step size 10 nt). Dashed vertical lines represent the predicted borders between IGR and the p200 and p110 ORFs.

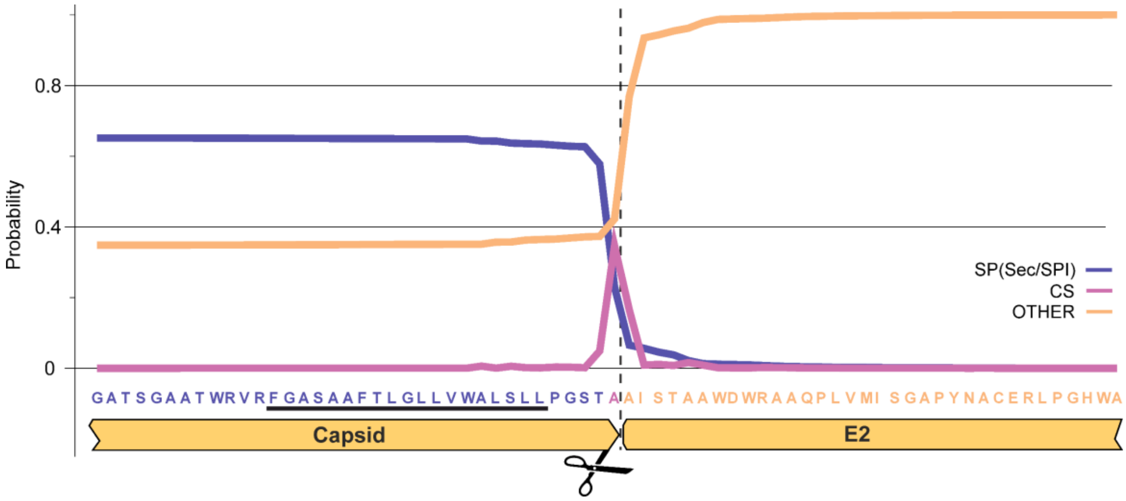


**Fig. S5.** Prediction of the signal peptidase cleavage site between capsid protein and E2 within the structural polyprotein p110 of rustrela virus. The probability for a signal peptidase recognition site (SP; blue), the signal peptidase cleavage site (CS; violet) and neither of both (OTHER; orange) is depicted. Prediction was carried out using the software package SignalIP5.0. The predicted transmembrane region in the capsid protein is highlighted by a black underline and the predicted SP cleavage site is shown as dashed line.

**Table S1**. Immunohistochemical markers and applications

| Marker | Antibody and dilution factor | Antigen Retrieval | Secondary reagents |
|---|---|---|---|
| **Active caspase 3** | Mouse anti-active Caspase 3 (Cell signalling, Leiden, The Netherlands), 1:400 overnight | HIER, Citrate buffer pH 6.0, for 20 min | secondary biotinylated goat anti-mouse antibody (Vector Laboratories, Burlingame, CA, USA), 1:200, 30 min; ABC Kit Vectastain (PK 6100, Burlingame, CA, USA), 30 min |
| **CD79a** | Mouse anti-CD79A (clone HM57) monoclonal, (LifeSpan BioSciences, Seattle, WA, USA), 1:50, overnight | HIER, 10mM Tris/ 1mM EDTA buffer pH 9.0, 20 min | Dako EnVision+ System- HRP Labelled Polymer Anti-mouse (Dako, Carpinteria, CA, USA), 30 min |
| **CD3** | Rabbit anti-CD3 polyclonal (Dako), 1:100, overnight | HIER, 10mM Tris/ 1mM EDTA buffer pH 9.0, 20 min | Dako EnVision+ System- HRP Labelled Polymer Anti-rabbit, 30 min |
| **Iba-1** | Rabbit anti-Iba1 (FUJIFILM Cellular Dynamics, Madison, WI, USA), 1:800, overnight | HIER, Citrate buffer pH 6.0, for 20 min | Dako EnVision+ System- HRP Labelled Polymer Anti-rabbit, 30 min |
| **GFAP** | Rabbit anti-GFAP (Abcam, Cambridge, UK), 1:200, overnight | HIER, Citrate buffer pH 6.0, for 20 min | Dako EnVision+ System- HRP Labelled Polymer Anti-rabbit, 30 min |

HIER, heat-induced epitope retrieval; HRP, horseradish peroxidase
